# Characterization of a recombinant plastid translocon co-chaperone

**DOI:** 10.1101/2021.06.07.447357

**Authors:** Salima Sarah Nurmohamed, Kenton Ko

**Affiliations:** Department of Biology, Queen’s University, Kingston, Ontario, Canada K7L 3N6; Department of Biochemistry, Queen’s University, Kingston, Ontario, Canada K7L 3N6; Department of Biochemistry, Sanger Building, University of Cambridge, Cambridge CB2 1GA UK

**Keywords:** Recombinant Tic40, Temperature effects, Function, Complexes

## Abstract

The transport of proteins into plastids is dependent on how the different translocon components work together. One translocon component, Tic40, possesses functional features found in other proteins such as co-chaperones and may thus work similarly in this capacity. To enhance the understanding of Tic40’s mode of operation, a more basic characterization of the protein itself is required. This study was designed largely to examine the properties of recombinant Tic40 without the predicted transmembrane region. The properties of Tic40 revealed in this report are: 1) In plants, overall Tic40 levels can adjust to temperature; 2) Recombinant Tic40 proteins appear to display some level of functionality in the soluble form; and 3) Outside of a membranous context, the formation of complexes involving recombinant Tic40 can be influenced by temperature. The potential significance of these basic properties is discussed.

## 1. Introduction

The population of pre-proteins being acquired by plastids at any one instance is extensive, structurally diverse, and continually adapting to the plant cell’s needs. Unexpected alterations, such as the change in protein structure, can also come into play when translocating pre-proteins are exposed to different temperature conditions like heat or cold. A responsive protein transport process must therefore be maintained by plastids to accommodate all situations, either immediately or for longer periods. The importance of being responsive to the protein transport needs of a plant cell has been demonstrated for plant mitochondria [1–4]; therefore, plastids will likely be responsive for the same reasons [5–8].

There are currently several adaptive mechanisms of the plastid protein transport machinery (translocon) under investigation, such as different receptors, alternate pathways, stoichiometry, and post-translational modifications [5–10]. Another potential adaptation mechanism may involve chaperones and co-chaperones associated with plastids. This possibility is based partly on how chaperone/co-chaperone systems in other cellular compartments work. For various purposes, chaperone/co-chaperone complexes are commonly involved with proteins considered aberrant, misfolded, unfolded or the like [9, 10]. For similar needs, such complexes are also involved in protein delivery and stress response activities, as examples. In the plastid translocon, components containing domains found in other chaperones or co-chaperones have been characterized, such as for Tic40 [11–13. Tic40 contains two potential co-chaperone features, a putative Sti1p/Hop and Hip domain (heat-shock protein interacting co-chaperones) and a weak version of tetratricopeptide repeats (TPR) [11, 12]. There are also defined chaperones and co-chaperones working with the plastid translocon, the Hsp70, Hsp90 and 93 proteins associated with different sites of the plastid envelope (from the cytoplasmic face of the outer membrane to the inner membrane) (see reviews [3, 7]). Proteomic data compiled for plastids indicate that large families of co-chaperone-like proteins exist (e.g., [14]). These proteins may also be participants of the plastid protein transport process. Chaperones and co-chaperones may thus operate in several ways to help the plastid translocon cope with the diversity of translocating proteins.

The Tic40 co-chaperone is a relatively abundant component of the plastid translocon. Some of the evidence from parallel lines of investigation suggests that the role of Tic40 is likely to be flexible, as opposed to an uncompromising central role [see reviews 5.6 for compilations]. Such findings suggest that Tic40 may be involved in some aspect of modulation in the translocon [15, 16] and that modulation of Tic40 stoichiometry may potentially influence how other translocon components work [15, 17]. In yeast, Tic40 was found to display an affinity to mitochondrial translocon components believed to partake in regulatory activities such as dimerization and co-chaperones [15]. Tic40 also exhibits the ability to work potentially as different configurations and associations [9, 10, 12, 13, 15, 16]. Tic40 was shown to stimulate two different activities during plastid protein import, transit-peptide release from Tic110 when recruited by Tic110 and ATP hydrolysis when associated with Hsp93 [9, 10, 18]. Tic40 also appears to assemble into the translocon at a stage much later than the other core components [19].

Although the features revealed so far provide insights into Tic40’s mode of operation, more basic characterization of the protein itself is important to understand its function. This study was thus designed solely to examine the various properties of recombinant Tic40 without the predicted transmembrane region. The main objective of the experiments is focused on how Tic40 works as a recombinant protein and how it responds to stress.

## 2. Materials and Methods

### 2.1. Plant material and procedures for the plant–based experiments

Plants were propagated at 21°C with a 16:8 h light:dark photoperiod (fluorescent and incandescent lighting). Specific conditions are noted when applicable. The production of transgenic *Arabidopsis thaliana* (cv. WS) plants used here was reported previously [16]. Chloroplasts were prepared from seedlings grown on media plates as described [20, 21]. Chlorophyll determinations were performed as reported by Porra et al. [22]. Total cellular protein extracts were prepared for the various indicated tissues by grinding fresh samples in pre-chilled extraction buffer as previously described [23]. The extracts were quantitated and normalized using Bradford protein assays before use. For comparative studies, equal amounts of samples, typically representing 0.5 μg total protein, were loaded per lane in the protein gels shown. Standard immuno-blotting protocols were used to assess the various extracts. Immunoreactive bands were scanned, quantitated, and assessed relative to appropriate internal references when necessary. The use of antibodies (for the various components) was pre-determined by titration assays in previous work. Control experiments were performed to determine the linearity of band signals for the antibodies used and the proteins being assessed. Scans were conducted for each replicate using non-saturated versions of the results presented.

### 2.2. Materials and procedures for the analysis of recombinant Tic40

Recombinant proteins were produced using *Escherichia coli* BL21(DE3) or JM109 (DE3) cells. The Tic40 cDNA sequences were isolated from *Brassica napus* cv. Topas (previously named *bnToc36B*, accession number X79091) [24, 25, 16]. Tic40 represents the full-length protein and Tic40s represents a shorter form (lacking the N-terminus region that encompasses the predicted transmembrane region). BnToc36B was originally isolated as a Tic40s form. Carboxyl-terminal histidine or N-terminal glutathione-S-transferase fusion tags were added using the respective plasmids pET21b (Novagen, Madison, Wisconsin, USA) and pGEX-4T3 (Amersham Biosciences, PQ, Canada). To add both tags, the pGEX-4T3 vector was modified to include an additional C-terminal 6XHis tag; the resultant plasmid was named pGEXCHIS. Such fusions have had no observable impact on the function or transport of the protein [26, 15]. Tic40s forms could also be generated for *R. communis* (designated as *rcTic40,* accession number DQ4735800) and *A. thaliana* (designated *atTic40*, accession number AY093010) using a unique *Bst*EII site in the cDNA sequence. Although not shown or reported in this study, Tic40s forms from these two other species behave in the manner as that observed for *B. napus* Tic40s. Proteins were purified using the appropriate affinity chromatography system. Cells were grown at 37°C in ampicillin-containing Terrific Broth (Bioshop Inc., Burlington, ON, Canada). Cells were induced using 0.8 mM isopropyl-β-D-1-thiogalactopyranoside (IPTG) at an OD_600_ of 0.8 for 4 hours. Cells were harvested in buffer A ((50 mM Na_2_HPO_4_, pH 8.0, 300 mM NaCl, 5 mM imidazole, 1 mM phenylmethane sulfonyl fluoride, 10 mM dithiotheritol and 0.1 mg ml^−1^ DNAse I), frozen and stored at −20°C. To purify the Tic40s protein, the cells thawed and then lysed in buffer A using a cell disrupter (Cell Disruption System, UK) at 25 kpsi. His-tagged proteins were recovered from the clarified lysate using a drip column containing a 5.0 ml bed volume of Ni-NTA resin (Qiagen, Mississauga, ON, Canada). After extensive washing with buffer A, bound proteins were eluted with a step gradient of increasing imidazole concentrations. Select fractions were concentrated and further purified using a HiLoad 16/60 G-200 gel filtration column on the AKTA FPLC system (Amersham Biosciences, Quebec, Canada). Peak fractions were pooled, concentrated to approximately 10 mg ml^−1^ in buffer A using a Centricon 10 kDa concentrator (Millipore, Etobicoke, ON, Canada), dialyzed extensively with 5 mM Na_2_HPO_4_ (pH 7.0), and stored at −80°C until use.

Blue native gel electrophoresis was carried out as described [27]. The visualization of bands representing complexes and monomeric counterparts was carried out using standard immunoblotting techniques.

Dynamic light scattering (DLS) (Dynapro light scatterer, DynaPro, UK) experiments were carried out at sequential temperatures of 4, 16, 24, and 40°C. Purified recombinant Tic40 proteins were prepared in phosphate buffer at a concentration of 10 mg ml^−1^. The data were modeled and analyzed using the accompanying software (DynaPro, UK).

Secondary structure analysis was performed on the 31 kDa recombinant Tic40 form by circular dichroism (CD) via a rapid scanning monochrometer (OLIS RSM 1000). Purified proteins were adjusted to 0.1, 0.2, and 0.3 mg ml^−1^ in 10 mM Na_2_PO_4_ (pH 7.0). Eight CD scans were performed at 25°C for each concentration using a 0.1 mm path length cuvette. The baseline was corrected by subtracting the spectra measured under identical conditions for the remaining buffer after dialysis. Secondary structure percentage was calculated using CDNN software [28].

Samples used in analytical ultracentrifugation (AUC) were dialyzed extensively against phosphate buffer (50 mM Na_2_PO_4_, pH 7.0, 150 mM NaCl). The dialysate of each sample was used for making protein dilutions and for the reference solution. All AUC experiments were performed at 20°C. Sedimentation equilibrium analysis was performed using a Beckman XL-I analytical ultracentrifuge with a four-hole An-60Ti rotor with 6-sector Epon charcoal centerpieces. Data were collected at 280 nm at three rotor speeds (15,000, 25,000, and 60,000 xg) and at three protein concentrations (0.1, 0.3, and 0.5 mg ml^−1^). Absorbance measurements were taken at 0.002 cm radial steps and averaged over 10 observations. Data sets were collected after equilibrium was obtained, as judged by the successive overlay of scans at 2-hour intervals. The partial specific volume and solution density were calculated by SEDNTERP (version 1.05, John Philo, 2000). The molecular weight for both a single ideal species and a monomer-trimer interaction were obtained via XL-A/XL-I data analysis software (version 6.03, Bechman/Microcal). The sedimentation velocity analyses were carried out at 20°C in cells containing double-sector Epon charcoal centerpieces. Velocity runs were conducted at 60,000 rpm by use of Rayleigh interference optics. Interference scans were taken at intervals of 1 min for 400 scans. Best-fit profiles according to the continuous distribution *c*(*S*) Lamm equation model from SEDFIT [29] were overlaid on the experimental data by use of every second scan.

Size-exclusion chromatography was performed with recombinant Tic40 proteins. Bacterial cells were grown at 37°C and induced with IPTG (at 0.4 OD_600_) for an additional 1.5 h. Cells were harvested, frozen, and then re-suspended in phosphate buffer. Lysates were prepared by sonication and then clarified by centrifugation at 20,000xg for 45 minutes. Clarified lysates were filtered, concentrated (to1-2 ml), and clarified before use. The samples contained approximately 20 mg of total protein and were loaded at a flow rate of 0.5 ml min^−1^ onto a Superdex G-200 (16/60) size exclusion column using the AKTA FPLC system (Amersham Biosciences, Quebec, Canada). Three-ml fractions were collected. The column was calibrated using standard molecular weight markers. The markers were run individually using a flow rate of 0.5 mg ml^−1^ (catalase, 232 kDa; aldolase,158 kDa; bovine serum albumin, 67 kDa; ovalbumin, 43 kDa; chymotrypsinogen A, 25 kDa; and ribonuclease A, 13.7 kDa). The void volume was calculated using blue dextran (2,000 kDa).

For the chromatography experiments, protein gel loadings were based on equal volumes. The volume used for a particular experiment was determined after assessing the protein content of each fraction. The anti-Tic40 antibodies used were equally immunoreactive for all forms, whether made in bacteria or plants [15, 16, 25]. Control experiments were performed to determine the linearity of band signals for the anti-Tic40 antibodies used and the proteins being assessed.

## 3. Results and discussion

### 3.1. Analysis of Tic40 levels in plants treated with different temperatures

Chaperones and co-chaperones are commonly defined by their stress-responsiveness and their involvement in stress-related activities. Tic40 was previously observed to fluctuate in two ways during plantlet and seed development, as different forms and in their relative abundance (for *Pisum sativum* and *B. napus*) [15]. This aspect of Tic40 may represent a response to the changes in the pre-protein population and the accompanying demands placed on the plastid translocon. Although these fluctuations occurred relatively fast, hourly for the plantlets, longer-term genetic programs may still govern this response. We therefore needed to see if Tic40 as a co-chaperone could also respond to externally applied stress in shorter time frame. Heat and cold are known to influence cellular activities, including protein transport processes, and were thus selected as the two conditions to test. *Arabidopsis* seedlings were subjected to short-term heat shock (4 hours at 37°C) and long-term cold acclimation (growth at 4°C) to look for a Tic40 response in mature leaves (Figures 1a and b). Short-term heat shock was applied to plants grown at 21 and 27°C. The results show that plants respond by increasing Tic40 levels (relative to Tic110) in all three cases (ANOVA for 21 to 37°C, F=13.37, P=0.021; for 27 to 37°C, F=71.03, P=0.001; for 4°C, F=101.46, P=0.0005). In all three cases, the increases were statistically significant. For comparative purposes, changes in stoichiometry between Tic40 and Tic110 are represented as normalized ratios relative to the wild-type samples. The “wild-type” ratios are set and defined as 1 for each of the experiments presented. Although Tic40 levels showed stress-responsiveness, the accumulation of the plastid proteins, Rbcs, Oee1 and Lhcb, was generally similar to the unstressed plants (Figure 1a). Import appears to be sustained at a level comparable to the wild-type plants during the treatment period. The levels of other translocon components were also similar to the wild-type plants (Figure 1a). To see how the actual plastid translocon reacts to changes in temperature, we opted to assess responses at the protein level only, rather than at the transcript level. These results indicate that Tic40, like other chaperones and co-chaperones, is responsive to stress by changing steady-state levels. Although Tic40 displays a common co-chaperone feature, the reason for altering steady-state Tic40 levels during stress is yet to be studied in further detail.

**FIGURE 1.**
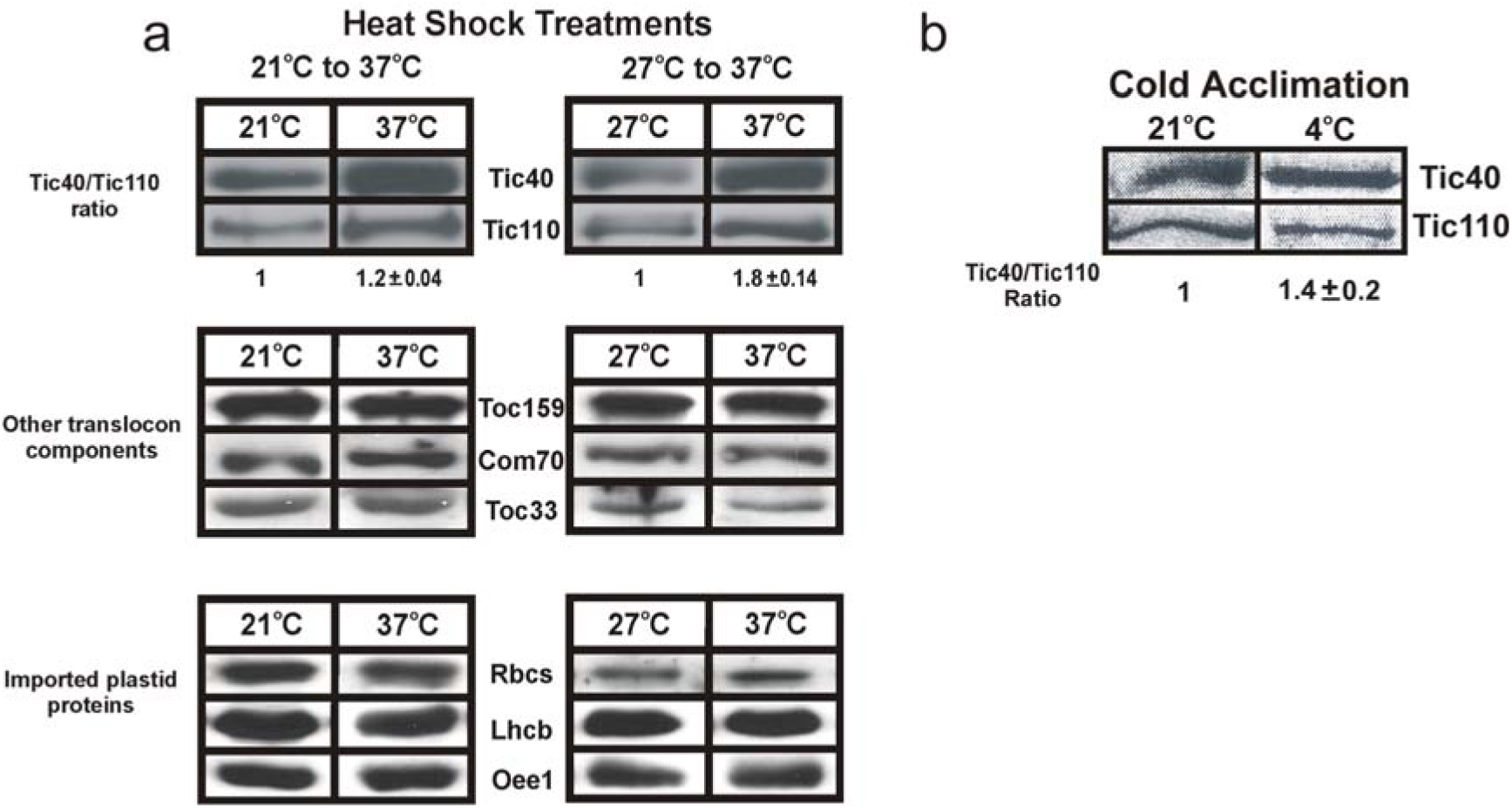
Tic40 displays stress-responsiveness in plants. (a) Immunoblot analysis of Tic40 levels in heat-treated *Arabidopsis* plants. The treatment schemes are indicated. Tic110 was analyzed for the same plastid samples and presented as a relative reference point. Changes in stoichiometry between Tic40 and Tic110 are represented as normalized ratios relative to the wild type (n=3, ± SD). The wild-type ratio is defined as 1 (set at 1). For comparison, the same samples were also analyzed with antibodies against other translocon components (Toc159, Com70, and Toc33) and imported plastid proteins (Rbcs (ribulose-1, 5-bisphosphate carboxylase in the stroma), Lhcb (chlorophyll *a/b* binding protein in the thylakoid membrane), and Oee1 (the 33 kDa oxygen-evolving subunit in the thylakoid lumen)). Equal amounts of total protein were loaded in each lane. Representative profiles are presented. (b) Immunoblot analysis of Tic40 levels in cold-acclimated *Arabidopsis* plants. Plants were grown at 21°C and 4°C for this comparison. As in panel A, Tic110 was analyzed for the same samples and presented as a relative reference point. Changes in stoichiometry between Tic40 and Tic110 are represented as ratios (n=3, ± SD). Representative profiles are presented.

Another indication of a potential link between Tic40 levels and stress-related activities can be observed when transgenic *Arabidopsis* seedlings were exposed to heat (37°C) for 4-6 days (Figure 2). Analysis was focused on two groups of transgenic plants, designated “Low” and “High” Tic40 levels. Relative to the wild-type plants grown under the same conditions, the Tic40 levels were suppressed approximately 3 times in the “Low” lines and elevated about 6 to 10 times in the “High” lines. The level of heat tolerance observed generally reflected the Tic40 levels (relative to wild type). Generally, tolerance to heat was slightly higher in seedlings with elevated Tic40 levels (“High” lines) and lower in plants with suppressed levels (“Low” lines). For the Tic40 suppressed lines (called “Low” lines), 258 out of the 275 seedlings assessed exhibited visible signs of damage caused by exposure to 37°C. In this study, heat tolerance is defined qualitatively as the number of days before visible signs of damage appear, e.g., wilting, decoloration, necrosis, and death. Growth impairment was not considered in these observations. Although many other factors are involved in heat tolerance, Tic40, like other co-chaperones, appears to be one possible participant. It is important to note that all plant lines grew well under normal conditions. There were no visible signs of unhealthy phenotypes.

**FIGURE 2.**
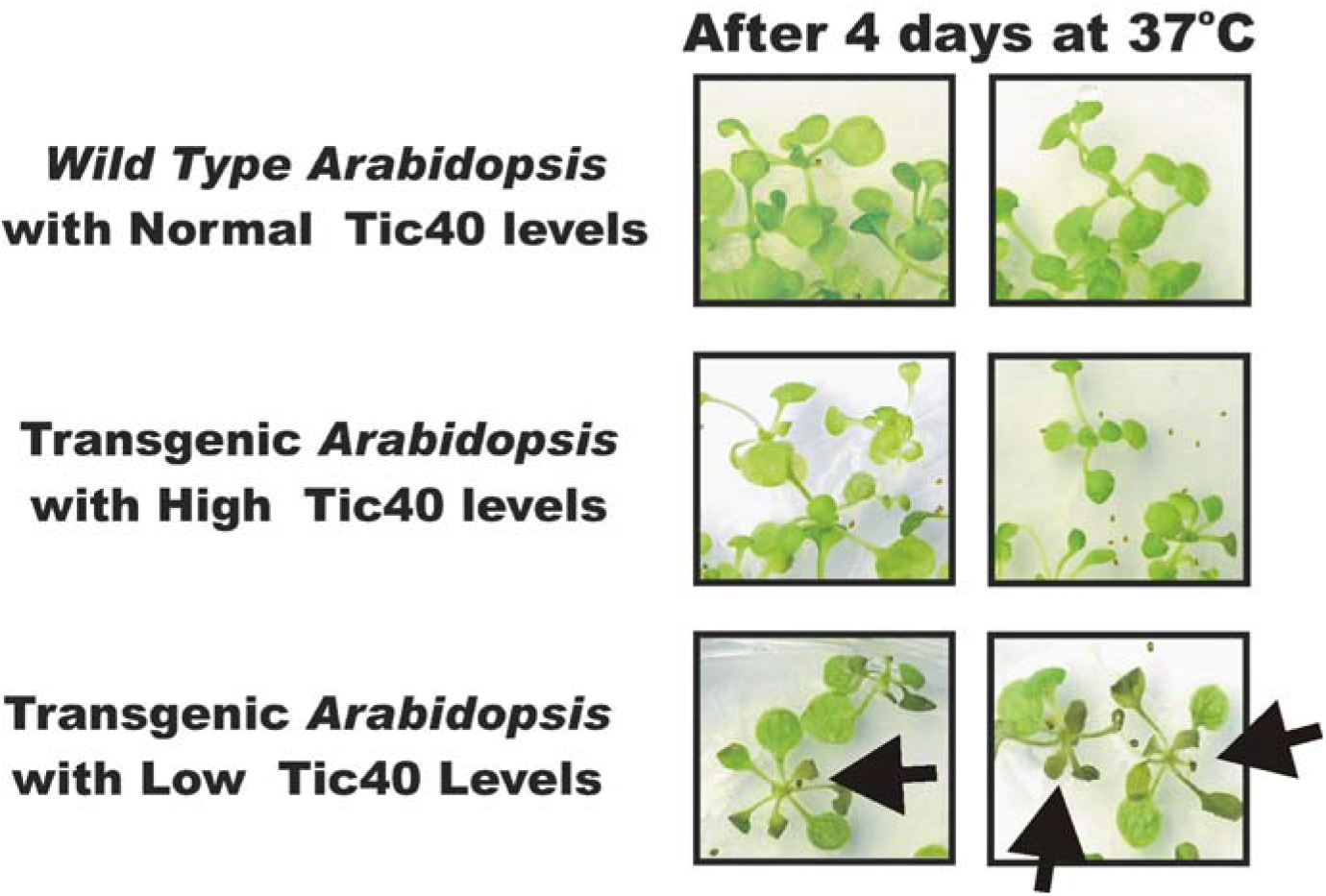
Transgenic *Arabidopsis* plants (from H1-3 and L1-3) were subjected to heat (37°C) for up to 4 days and documented. Relative to the control wild-type plants, transgenic plants with excessive production of Tic40 exhibited more tolerance to heat whereas plants with depressed production displayed less tolerance (highlighted with arrows in the latter case). The experiments were carried out three separate times. The lines used are the same those used in Figure 1c. The transgenic plants with low Tic40 levels showed signs of heat damage in 258 of the 275 seedlings assessed. Representative samples are shown.

### 3.2 Analysis of recombinant Tic40 in cell extracts and as purified proteins

The basic co-chaperone characteristics displayed above suggest that Tic40 levels may reflect the protein’s ability to form Tic40-containing complexes since most of the other translocon components appear to be relatively unresponsive to stress. The experiments in this section were thus designed solely to find out if recombinant Tic40 proteins possess such a property. First, we used total cell extracts prepared from Tic40-expressing bacterial strains to see if there was an indication that recombinant Tic40 formed complexes with itself, i.e. a population of complexes containing more than one recombinant Tic40 protein in whole bacterial cell extracts. These complexes can be created directly, indirectly, or both, and may contain other bacterial proteins. The extracts were analyzed by size-exclusion chromatography and immunoblotting (Figure 3a). The recombinant Tic40 proteins used are without the predicted transmembrane region (designated hereon in as Tic40s) and are His-tagged for identification purposes. All co-chaperone signature domains remain intact in Tic40s. Recombinant Tic40s proteins do retain some form of functionality, as shown in a number of other studies [15, 16, 18]. We also incorporated two other tools to help in the assessment: 1) We used the smaller recombinant Tic40s proteins that arise during expression in bacteria as indicators of composition, and 2) We used GST fusion tags (N-terminus) to create deliberate shifts to portions of the population of recombinant Tic40s-containing complexes. If recombinant Tic40s can form complexes with itself, the gel filtration pattern would shift accordingly when fused to large GST tags and some of the smaller Tic40s proteins (40 and/or 31 kDa) present in the complexes may shift along with the larger GST-tagged Tic40s. The 31 kDa form arose from a start site downstream (see below for more details). If Tic40s proteins were strictly monomeric in nature, the gel filtration profiles would display independent shifts, i.e. the smaller Tic40s forms would remain in the same fractions, separate from the GST-Tic40s fusion proteins. The resulting profiles appear to indicate that Tic40s is capable of forming a population of complexes containing more than one Tic40s protein and with different combinations of Tic40s forms (Figure 3). The fusion of GST tags created shifts in the population of Tic40s-containing complexes. The population of complexes that shifted contains more than one Tic40s protein, as revealed by the antibodies against the C-terminal His-tag. In the His-tagged Tic40s profile, the relative molecular masses for complexes eluting in fractions 21 and 25 were estimated to be around 180 and 66 kDa, respectively. In the GST- and His-tagged Tic40s profile, the relative molecular mass for complexes eluting in fractions 18 was estimated to be around 265 kDa. Tic40s proteins alone do not bind to the GST tag itself (Figure 3b). This was the case for both His-tagged and untagged Tic40s proteins. Interactions can also be observed to occur between GST-tagged Tic40s and His-tagged Tic40s when brought together and assessed by GST affinity chromatography (Figure 3c).

**FIGURE 3.**
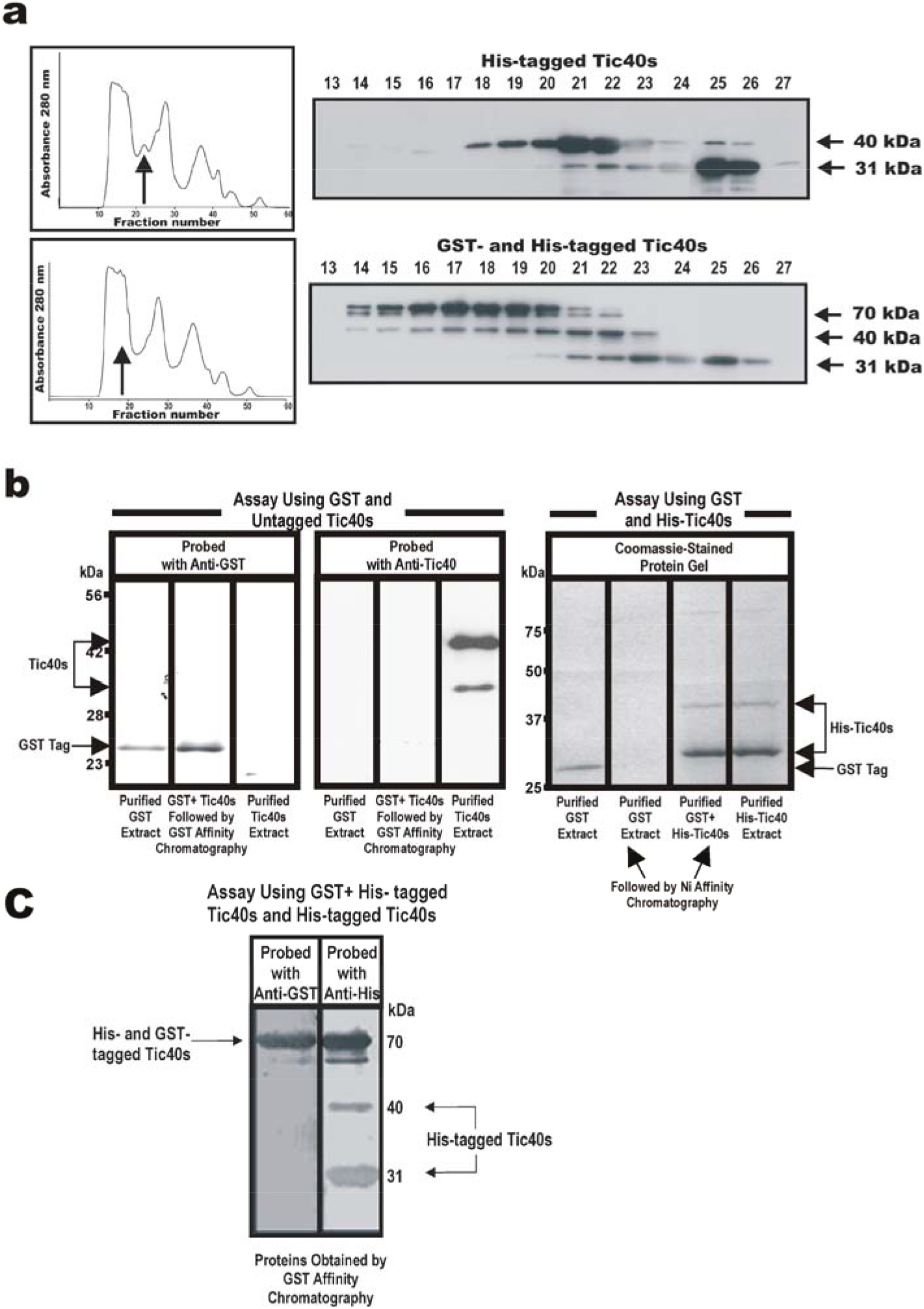
Gel filtration profiles of recombinant Tic40s-containing complexes produced in bacterial cells. (a) Profiles of complexes produced with carboxyl terminus histidine-tagged Tic40s (top half) and with the GST-Tic40s-His tag construct (lower half). For SDS-PAGE, the same volume was used per lane. Immunoblots were analyzed with antibodies against the His tag. Corresponding chromatograms are provided to the left of the immunoblots. The arrows in each of the chromatograms mark the shifting event discussed in the text (the top arrow indicates the position of the Tic40s-containing complexes without the GST tag and the bottom arrow indicates the position of the Tic40s-containing complexes with the GST tag). The relative molecular masses of the different Tic40 bands are indicated to the right of the immunoblots. Each immunoblot is derived from two simultaneously run SDS protein gels (from lanes 13-24 and 25-27). A representative profile is presented in this panel. In the His-tagged Tic40s profile, the relative molecular masses for complexes eluting in fractions 21 and 25 were estimated to be 180 and 66 kDa, respectively. In the GST- and His-tagged Tic40s profile, the relative molecular mass for complexes eluting in fraction 18 was estimated to be 265 kDa. (b) The degree of interaction between the two protein tags used in panel A, GST and His, was tested and presented here. The left set of lanes represents an assay between GST and untagged Tic40s. Both proteins were purified and mixed before GST affinity chromatography. The resulting samples were then assessed immunologically as indicated. The right set of lanes represents an assay between GST and His-tagged Tic40s. Both proteins were purified and mixed before Ni-affinity chromatography. The resulting samples were then assessed by SDS-PAGE. The assays on the left represent immunoblots from 12% SDS-PAGE gels and the assays on the right represent patterns from 10% SDS-PAGE gels. (c) Interactions between GST-tagged Tic40s and His-tagged Tic40s were also observed in a separate assay. Both of the tagged Tic40s proteins were brought together and assessed using GST affinity chromatography. The resulting samples were analyzed by immunoblotting as indicated.

Next, we proceeded to determine if purified recombinant Tic40s was capable of forming complexes, as opposed to the complexes existing in whole bacterial cell extracts used in the above examination. Histidine-tagged Tic40s proteins, purified by affinity chromatography and gel filtration, were analyzed immunologically (Figure 4). The immunoblots provide another qualitative indication that Tic40s was capable of forming complexes. Even though the proteins were purified through a two-stage process, the 40 kDa Tic40s form co-eluted with the smaller 31 kDa form. Complete separation of the 40 kDa Tic40s form into its own fraction was not entirely possible. The 31 kDa form, however, was separable in its own fraction. The identity of the 31kDa form was confirmed to be Tic40 by mass spectrometry and contained the co-chaperone signature domains present in the carboxyl segment of the protein (data not shown). Like the above outcomes, these results represent another qualitative indication that purified recombinant Tic40s may possess the ability to form complexes with itself.

**FIGURE 4.**
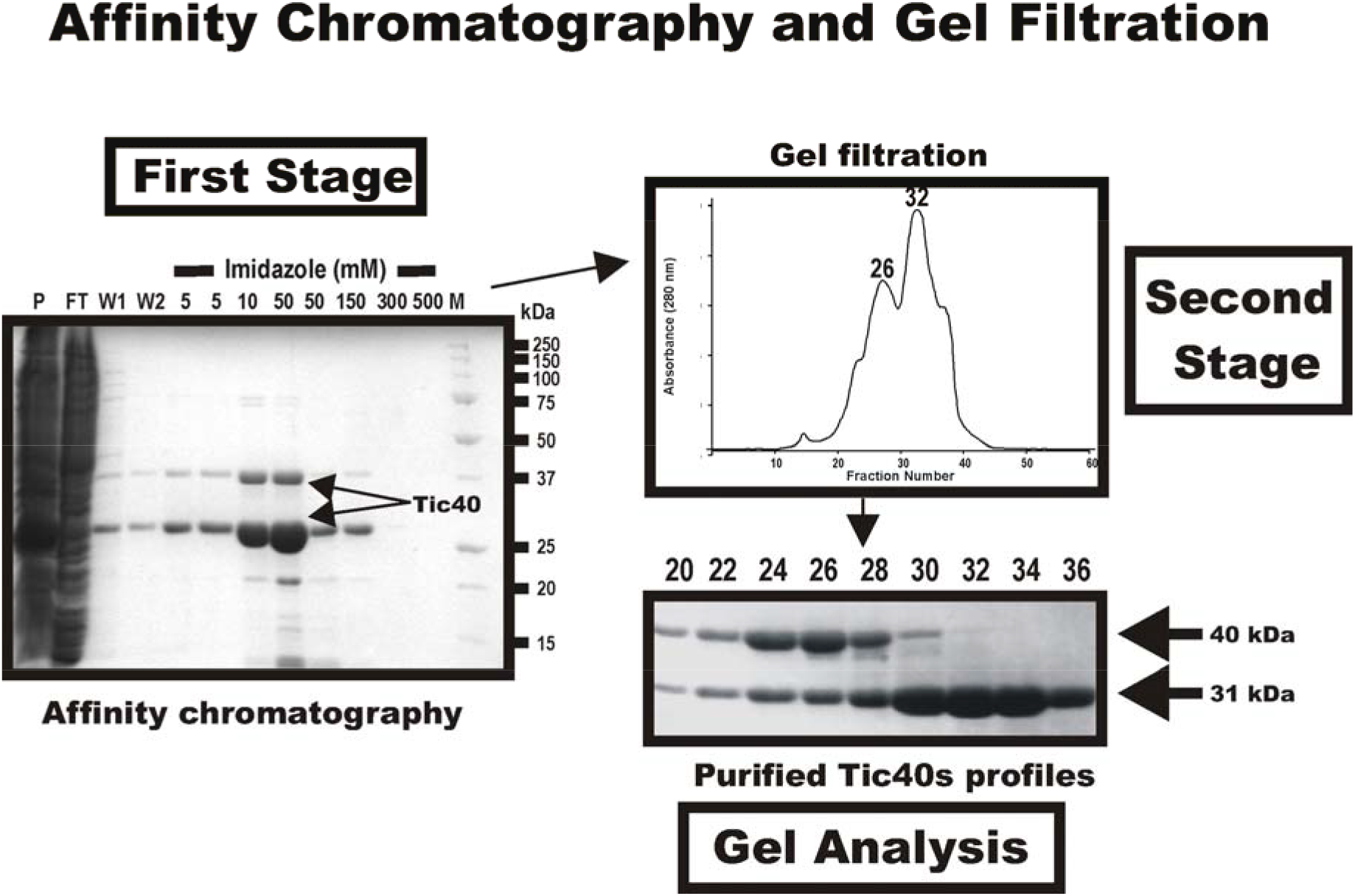
Characterization of complexes formed with purified recombinant Tic40s proteins. Affinity purification and gel filtration profiles of recombinant Tic40s proteins are presented. For the affinity purification profile of the first stage, the various fractions present in the Coomassie-stained SDS protein gel are: P, pellet; FT, flow-through; WT1 and WT2, washes 1 and 2; 5 to 500, step elutions at different imidazole concentrations; and M, marker. Fractions containing affinity-purified products were then subjected to gel filtration in the second stage. The resulting chromatogram is presented along with a Coomassie-stained protein gel of select fractions. For SDS-PAGE, the same volume was used per lane. The locations of fractions 26 and 32 are indicated on the chromatogram. The two Tic40s forms are marked with their corresponding relative molecular masses. All relative molecular masses are given in kDa. A representative profile is presented in this figure.

Blue native gel electrophoresis was also used as another qualitative approach to look solely for the presence of recombinant Tic40s-containing complexes and to see if there was an indication of temperature-responsiveness in the formation of these complexes (Figure 5). The bacterial strains analyzed were cultivated at 22 and 37°C. Separated complexes were then visualized immunologically. The results indicate the presence of Tic40s-containing complexes and that the proportion of Tic40s-containing complexes may be influenced by temperature (proportion of complex to total Tic40s in the sample). These bands or complexes (shown in Figure 5) are specific to the experimental strain and are not present in the control extracts.

**FIGURE 5.**
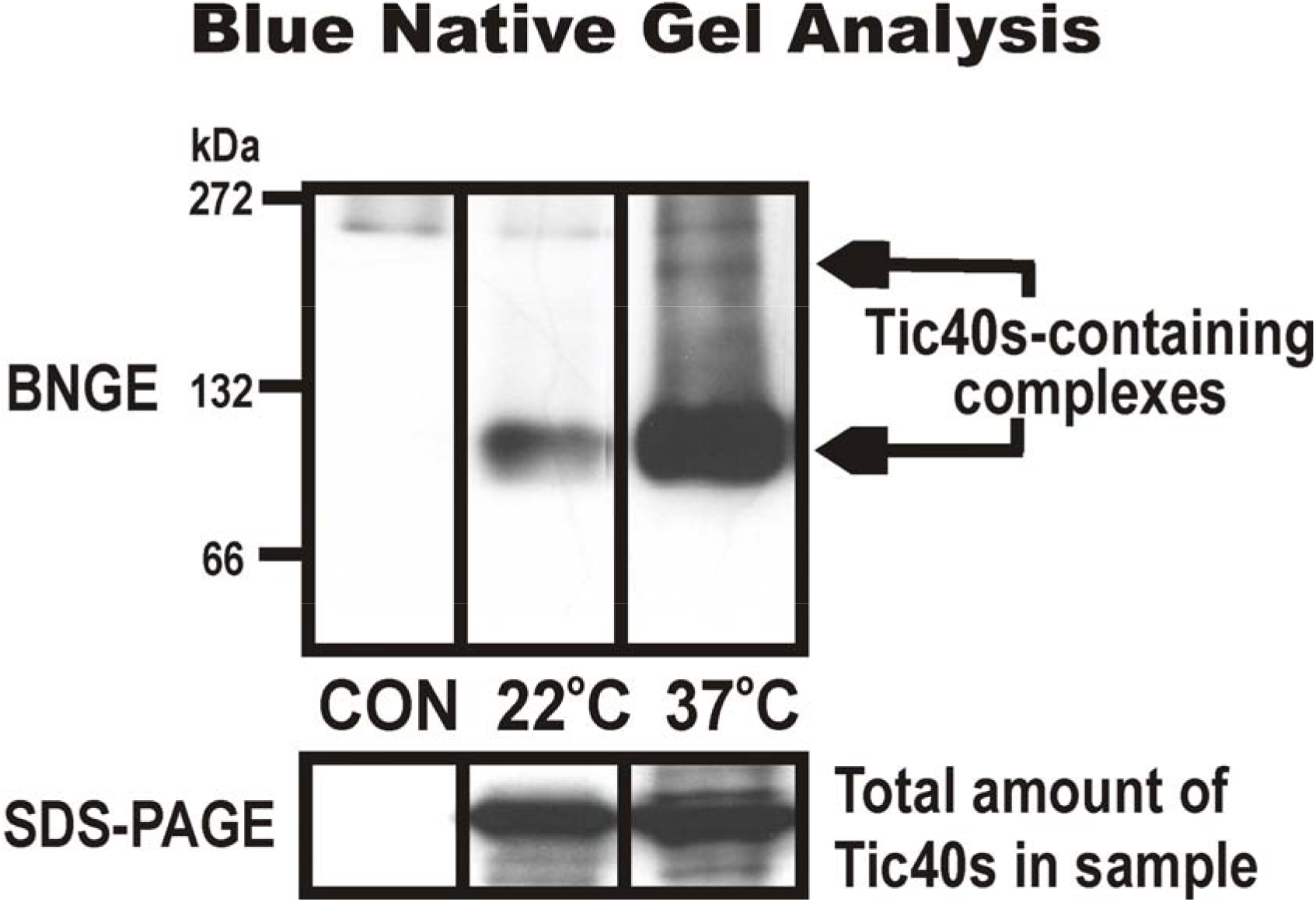
Analysis of recombinant Tic40s-containing complexes produced in bacteria using blue native gel electrophoresis (BNGE), SDS denaturing gel (SDS-PAGE), and immunoblotting. Both Tic40s-containing complexes and monomers are highlighted. Temperatures used in the experiments are indicated. The control extract is labeled as CON. Relative molecular masses are given in kDa.

To determine the functional status of the purified recombinant proteins, the secondary structure of Tic40s was first assessed by circular dichroism (CD) spectrometry (Figure 6). Since the 31 kDa form can be purified in its own fraction, we used this form for the analysis. The 31kDa form was confirmed to be Tic40 by mass spectrometry and contained the co-chaperone signature domains (data not shown). The CD spectrum indicates that the protein is well-ordered and possesses secondary structural features typical for a globular protein. We next used sedimentation equilibrium analytical ultracentrifugation (AUC) to further study the structural behavior of the complexes formed with purified recombinant Tic40s (fractions containing 40 and 31 kDa forms). By fitting the AUC data to a single-species model, the apparent weight-average molecular mass suggests that Tic40s self-associates in the native form (Figure. 7). This was further corroborated by molecular weight versus concentration analysis, which showed a concentration-dependent association of Tic40s (unpublished observations).

**FIGURE 6.**
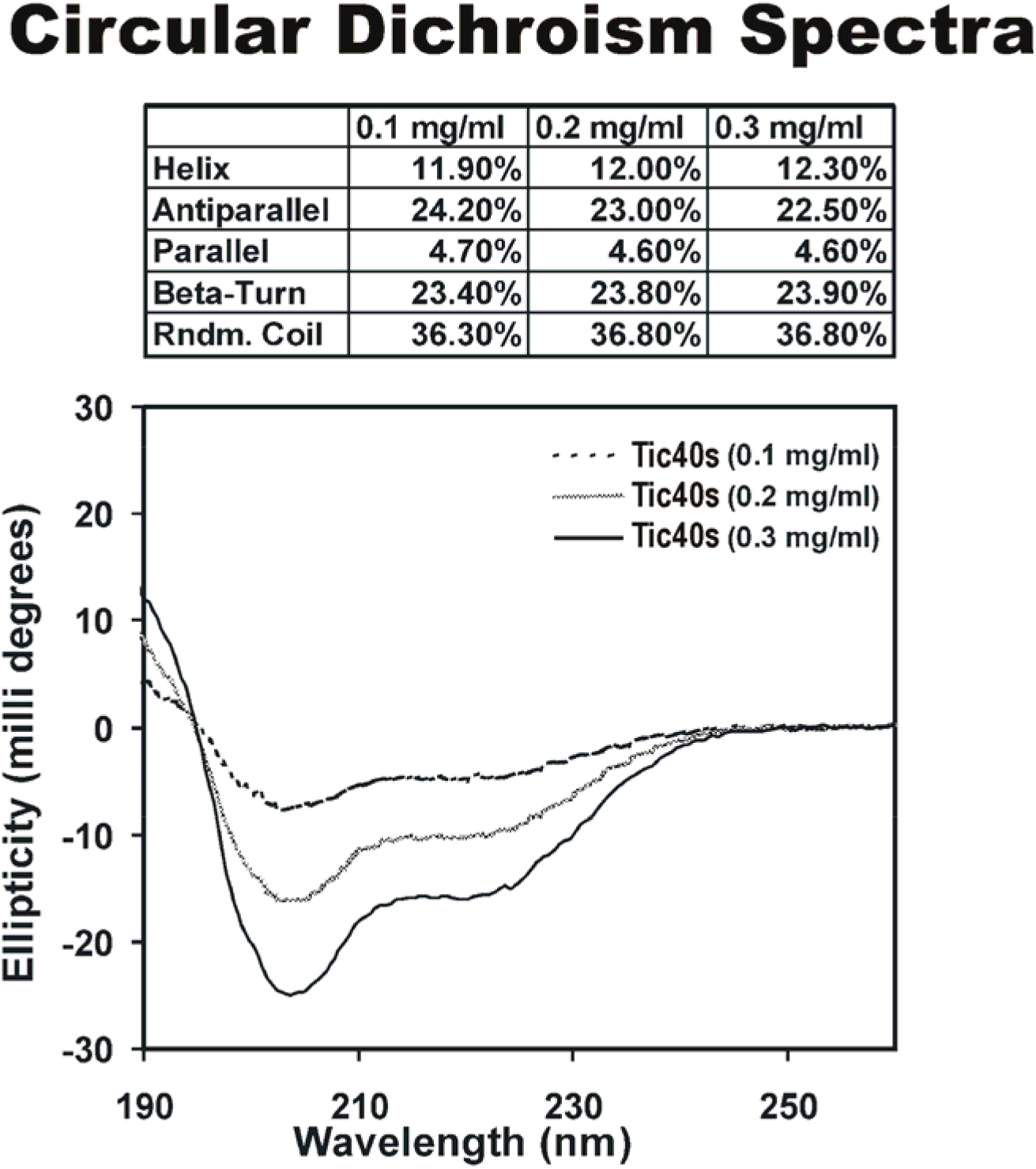
Circular dichroism spectrum for recombinant Tic40s proteins. The 31 kDa Tic40s was used at different concentrations. The analysis was conducted at 25°C.

**FIGURE 7.**
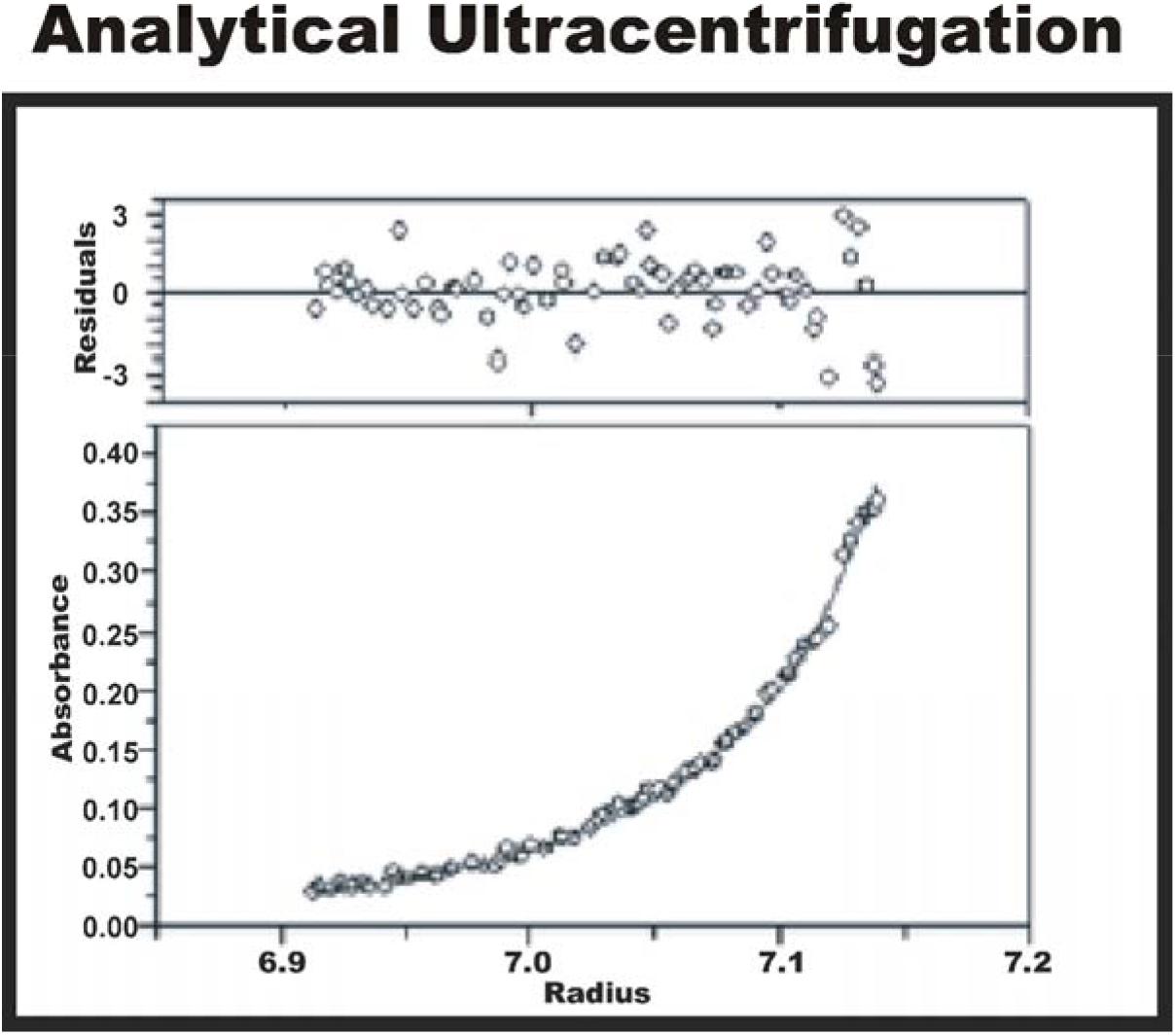
Assessment of complexes formed with recombinant Tic40s using analytical ultracentrifugation. Residual and absorbance plots (top and bottom panels, respectively) from a sedimentation equilibrium run of Tic40s are shown. The UV absorbance gradient (◊) in the centrifuge cell is shown. The data were fitted to a monomer-trimer equilibrium and the solid line denotes the fitted curve calculated from three rotor speeds (15,000, 25,000, and 60,000 xg, respectively) using 0.1, 0.3 and 0.5 mg ml^−1^ protein concentrations. Residuals show the difference in the fitted and experimental values as a function of radial position.

Sedimentation velocity experiments were also performed to assess the hydrodynamic properties of purified recombinant Tic40s (data not shown). Sedimentation profiles were evaluated by a continuous distribution *c*(*S*) Lamm equation model. Two peaks were revealed when the sedimentation coefficient distribution was determined, a major one at 2.13 S and a minor one at 5.19 S. These peaks corresponded to molecular masses of approximately 32,042 Da and 134,000 Da, respectively, indicating predominantly a monomer-tetramer type association.

We also analyzed various Tic40s fractions using dynamic light scattering (DLS). Fifty measurements were taken at temperatures of 4, 16, 24, and 40°C. The data for fractions containing both the 40 and 31 kDa Tic40s forms indicate that the proteins exist as a single species (Figure. 8) with approximately 12 % polydispersity. The corresponding hydrodynamic radius values (Rh) are approximately 25, 26, 29, and 48 nm. Based on the standard size versus weight relationship for globular proteins, the apparent molecular weight was approximately 70 kDa at 4°C and 110 kDa at 40°C. The DLS results for fractions containing only the 31 kDa Tic40s form indicate that this form exists in the dimeric state at lower temperatures and in a tetrameric state at higher temperatures (unpublished observations). These combined results indicate that the formation of complexes with purified recombinant Tic40s was possible and that their formation can be influenced by temperature. Larger Tic40s-containing oligomers tend to appear more when higher temperatures were used.

**FIGURE 8.**
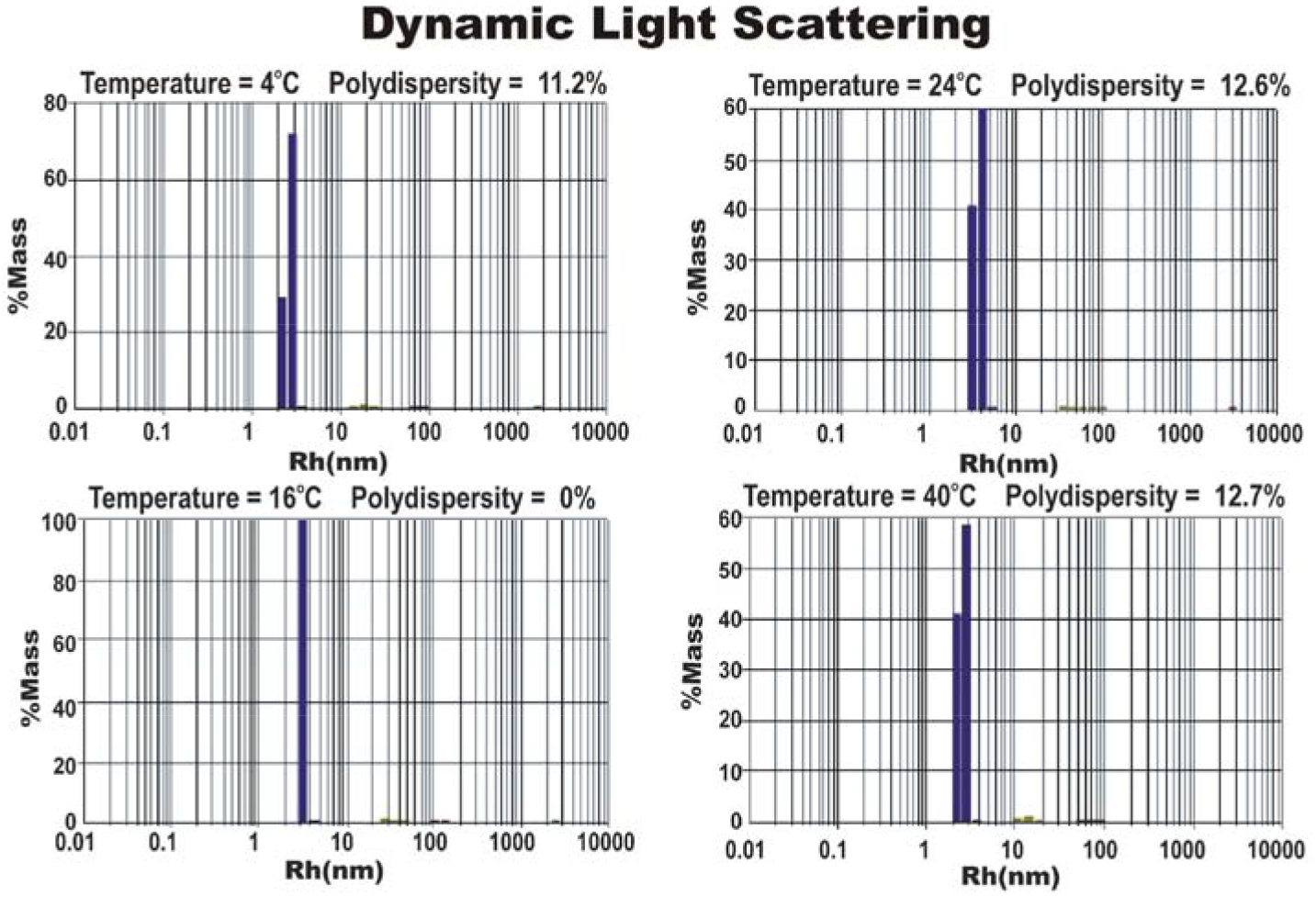
Dynamic light scattering patterns of recombinant Tic40s complexes at different temperatures. The level of polydispersity is given for each analysis.

## 4. Conclusion

Chaperones and co-chaperones exist throughout a cell and are involved in a variety of activities like protein delivery, regulatory functions, and stress response [9, 10]. Their roles in such activities are often flexible in nature, manifesting generally as transient interactions, responsiveness at several levels of the protein, and the ability to work as alternate forms. Like other protein transport systems, the plastid translocon also appears to involve components related to chaperones and co-chaperones. Tic40 is one such co-chaperone example [12]. Although the presence of potential co-chaperone features suggests that Tic40 is likely to work like co-chaperones, other basic properties related to the component’s co-chaperone definition needs to be assessed. This study was thus designed mainly to examine qualitatively other basic properties of Tic40 using recombinant proteins. The main characteristic revealed suggests that Tic40 possesses the potential to operate in a flexible manner.

The flexible nature of Tic40’s activities was observable in different contexts and at different levels. In whole plants, Tic40 responds to temperature by adjusting its level relative to the other core translocon components. Overall, there appears to be a need to adjust the plastid translocon in different situations, and one possible mechanism would likely be adjusting stoichiometry. Such a mechanism may be part of the reason for the slightly higher level of heat tolerance displayed by the transgenic *Arabidopsis* plants. Raising Tic40 levels may possibly help enhance the effectiveness of the plant’s stress coping mechanisms to a limited degree. Also, Tic40’s response to stress (temperature stress in this study) can be relatively swift, occurring within a 4-hour exposure to heat (in mature *Arabidopsis* leaves) or in hourly increments (previously observed in greening seedlings) [15]. Tic40’s response to temperature can even manifest at the level of the protein itself. The formation of Tic40s-containing complexes, either in association with other proteins or with itself, can be influenced to an extent by temperature. There is an indication that part of the response involves the engaging and disengaging of protein-protein interactions. The complexes of purified recombinant Tic40s themselves appear to possess the ability to change oligomeric status in different temperatures, for example, from a monomeric to a tetrameric state.

Transient interaction capabilities, modularity, and the ability to work as different forms are other features of Tic40 that may reflect flexibility as well. These qualities showed up in a number of experimental contexts. Although speculative at this point, these capabilities may be related to the adjusting of component levels such as that observed for Tic40 in plants. For instance, the increases in Tic40 levels in response to temperature change may reflect a mechanism for influencing how the plastid translocon works in different situations. Transient, modular capabilities were also evident in the formation of recombinant Tic40s complexes, for example, the presence of monomer-tetrameric states of oligomerization or complexes formed with different Tic40s forms. The same data also indicate that Tic40 is likely to be active and functional as different configurations.

Overall, the characteristics exhibited by Tic40s in this study indicate a substantial degree of flexibility in its capabilities and activities. These features mirror those established for other co-chaperones. Although most of the experiments conducted in this study were based on heterologous and *in vitro* approaches, the features revealed so far appear to reflect what Tic40 may be capable of doing, and hence how Tic40 may perform its role in the plastid translocon. This speculation extends from our preliminary work indicating that many of the properties revealed in this study appear to be in play in plants and in the membranous context of the plastid. Thus, the findings established in this study will help develop strategies for further investigations into Tic40’s mode of operation in the actual plastid translocon and how these Tic40 roles are carried out in plants.

## Acknowledgments

The authors thank Drs. Vinay Singh, Kim Munro and Zongchao Jia for their invaluable advice, training, and access to materials and equipment. The authors also thank Kelvin Chan and Lilian Lee for their assistance in conducting specific experiments while working on their own research projects. This work was supported by grants from the Natural Sciences and Engineering Research Council of Canada (K.K.), a Natural Sciences and Engineering Research Council of Canada Undergraduate Research Summer Award for her own work (to Lilian Lee), an Ontario Graduate Scholarship for his own work (to Kelvin Chan) and a Protein Function Discovery Program Fellowships from the Canadian Institutes for Health Research for this work (to S.N.).

## References

[1] R. Lister, O. Chew, M.-N. Lee J. L. Heazlewood, R. Clifton, K. L. Parker, A. H. Millar, and J. Whelan, “A transcriptomic and proteomic characterization of the *Arabidopsis* mitochondrial protein import apparatus and its response to mitochondrial dysfunction,” Plant Physiology, vol. 134, no. 2, pp. 777–789, 2004.

[2] N. L. Taylor, C. Rudhe, J. M. Hulett, T. Lithgow, E. Glaser, D. A. Day, A. H. Millar, and J. Whelan, “Environmental stresses inhibit and stimulate different protein import pathways in plant mitochondria,” FEBS Letters, vol. 547, no. 1-3, pp. 125–130, 2003.

[3] A. S. Ghifari, M. Gill-Hille, and M. W. Murcha “Plant mitochondrial protein import: the ins and outs,” Biochemical Journal, vol. 475, no. 13, pp. 2192–2208, 2018.

[4] M. W. Murcha, B. Kmiec, S Kubiszewski-Jakubiak, P.F. Teixeira, E. Glaser, and J. Whelan. “Protein import into plant mitochondria: signals, machinery, processing, and regulation,” Journal of Experimental Botany, vol. 65, no. 22, pp. 63016335, 2014.

[5] F. Kessler and D. Schnell, “The function and diversity of plastid protein import pathways: A multilane GTPase highway into plastids,” Traffic, vol. 7, no. 3, pp. 248–257, 2006.

[6] K. Ko, K. Chan, K. Karakasis, and B. Pedram, “Plastid protein delivery: coping with diversity,” Canadian Journal of Botany, vol. 84, no. 4, pp. 543–550, 2006.

[7] L.G.L. Richardson, D.J. Schnell. “Origins, function, and regulation of the TOC-TIC general protein import machinery of plastids,” Journal of Experimental Botany, vol. 71. No. 4, pp 1226–1238, 2020.

[8] B. Bölter. “En route into chloroplasts: preporteins’ way home”, Photosynthesis Research, vol. 138, pp. 263–275, 2018.

[9] A. J. Caplan, “What is a co-chaperone?” Cell Stress & Chaperones, vol. 8, no. 2, pp. 105–107, 2003.

[10] J. Frydman, “Folding of newly translated proteins *in vivo*: the role molecular chaperones,” Annual Review of Biochemistry, vol. 70, pp. 603–647, 2001.

[11] J. Bédard, K. Kubis, S. Bimanadham, and P. Jarvis, “Functional similarity between the chloroplast translocon component, Tic40, and the human co-chaperone, *Hip*,” The Journal of Biological Chemistry, vol. 282, no. 29, pp. 21404–21414, 2007.

[12] M.-L. Chou L. M. Fitzpatrick, S. H. Tu, G. Budziszewski, S. Potter-Lewis, M. Akita, J. Z. Levin, K. Keegstra, “Tic40, a membrane-anchored co-chaperone homolog in the chloroplast protein translocon,” The EMBO Journal, vol. 22, no. 12, pp. 2970–2980, 2003.

[13] T. Stahl, C. Glockmann, J. Soll, and L. Heins, “Tic40, a new ‘old’ subunit of the chloroplast protein import translocon,” The Journal of Biological Chemistry, vol. 274, no. 52, pp. 37467–37472, 1999.

[14] J. B. Peltier, J. Ytterberg, Q. Sun, and K. J. van Wijk, “New functions of the thylakoid membrane proteome of *Arabidopsis thaliana* revealed by a simple, fast, and versatile fractionation strategy,” The Journal of Biological Chemistry, vol. 279, no. 47, pp. 49367–49383, 2004.

[15] K. Ko, D. Taylor, P. Argenton, J. Innes, B. Pedram, F. Seibert, A. Granell, and Z. Ko, “Evidence that the plastid translocon Tic40 components possess modulating capabilities,” The Journal of Biological Chemistry, vol. 280, no. 1, pp. 215–224, 2005.

[16] K. Ko, S. Banerjee, J. Innes, D. Taylor, and Z. Ko, ‘The Tic40 translocon components exhibit preferential interactions with different forms of the Oee1 plastid protein precursor,” Functional Plant Biology, vol. 31, no. 3, pp. 285–294, 2004.

[17] S. Kovacheva, J. Bédard, R. Patel, P. Dudley, D. Twell, G. Ríos, C. Koncz, and P. Jarvis, “*In vivo* studies on the roles of Tic110, Tic40 and Hsp93 during chloroplast protein import,” The Plant Journal, vol. 41, no. 3, pp. 412–428, 2005.

[18] M.-L. Chou C.-C. Chu L.-J. Chen M. Akita, and H.-M. Li “Stimulation of transit-peptide release and ATP hydrolysis by a cochaperone during protein import into chloroplasts,” Journal of Cell Biology, vol. 175, no. 6, pp. 893–900, 2006.

[19] K.-Y. Chen and H.-M. Li “Precursor binding to an 880-kDa Toc complex as an early step during active import of protein into chloroplasts,” The Plant Journal, vol. 49, no. 1, pp. 149–158, 2007.

[20] H. Aronson and P. Jarvis, “A simple method for isolating import-competent *Arabidopsis* chloroplasts,” FEBS Letters, vol. 529, no. 2-3, pp. 215–220, 2002.

[21] L. Fitzpatrick and K. Keegstra, “A method for isolating a high yield of *Arabidopsis* chloroplasts capable of efficient import of precursor proteins,” The Plant Journal, vol. 27, no. 1, pp. 59–65, 2001.

[22] R. J. Porra, W. A. Thompson, and P. E. Kriedemann, “Determination of accurate coefficients and simultaneous equations for assaying chlorophyll *a* and *b* extracted with four different solvents: verification of the concentration of chlorophyll standards by atomic absorption spectroscopy,” Biochimica et Biophysica Acta, vol. 975, no. 3, pp. 384–394, 1989.

[23] W. C. Plaxton, “Molecular and immunological characterization of plastid and cytosolic pyruvate kinase isozymes from castor-oil-plant endosperm and leaf,” European Journal of Biochemistry, vol. 181, no. 2, pp. 443–451, 1989.

[24] K. Ko and Z. Ko, “Carboxyl-terminal sequences can influence the *in vitro* import and intraorganellar targeting of chloroplast protein precursors,” The Journal of Biological Chemistry, vol. 267, no. 20, pp. 13910–13916, 1992.

[25] K. Ko, D. Budd, C. Wu, F. Siebert, L. Kourtz, and Z. W. Ko, “Isolation and characterization of a cDNA clone encoding a member of the Com44/Cim44 envelope components of the chloroplast protein import apparatus,” The Journal of Biological Chemistry, vol. 270, no. 48, pp. 28601–28608, 1995.

[26] K. Ko and Z. Ko, “*In vitro* targeting of the Toc36 component of the chloroplast envelope protein import apparatus involves a complex set of information,” Biochimica et Biophysica Acta, vol. 1421, no. 1, pp. 198–206, 1999.

[27] L. G. J. Nijtmans, N. S. Henderson, and I. J. Holt, “Blue native electrophoresis to study mitochondrial and other protein complexes,” Methods, vol. 26, no. 4, pp. 327–334, 2002.

[28] G. Bohm, R. Muhr, and R. Jaenicke, “Quantitative analysis of protein far UV circular dichroism spectra by neural networks,” Protein Engineering, vol. 5, no. 3, pp. 191–195, 1992.

[29] P. Schuck, M. A. Perugini, N. R. Gonzales, G. J. Howlett, and D. Schubert, “Size-distribution analysis of proteins by analytical ultracentrifugation: strategies and application to model systems,” Biophysical Journal, vol. 82, no. 2, pp. 1096–1101, 2002.

